# Space-efficient method for high-throughput generation of uniform cell-laden hollow agarose microcapsules

**DOI:** 10.1101/2025.10.19.683215

**Authors:** Bin Xu, Sunwoo Kim, Kenji Hinode, Kazuki Hattori, Sadao Ota

**Affiliations:** Research Center for Advanced Science and Technology, The University of Tokyo, 4-6-1 Komaba, Meguro-ku, Tokyo 153-8904, Japan

**Author notes:** Co-first authors (Bin Xu, Sunwoo Kim).

## Abstract

Hollow hydrogel microcapsules isolate cells while allowing diffusion of nutrients and factors, supporting applications from single-cell analysis and clonal expansion to spheroid formation inside them. Conventional encapsulation relies on bulky instruments such as pumps and pressure controllers, hindering use in confined environments such as anaerobic chambers. Here we present a space-efficient method combining pump-free, −20 kPa negative-pressure droplet generation with particle-templated emulsification to produce millions of ∼70 µm hollow agarose microcapsules using only a sealed bottle and a vortex mixer within a 30 cm × 30 cm footprint. The system maintains ∼491 ± 2 drops/s and yields (2.26 ± 0.09) × 10^6^ capsules per batch with CV < 4% (N = 3). Operating entirely inside an anaerobic chamber, we encapsulated mouse gut bacteria and quantified population-level proliferation over four days. This space-efficient workflow enables high-throughput encapsulation under spatial constraints such as biosafety cabinets or anaerobic chambers, providing a practical foundation for downstream single-cell and clonal analyses.

## Introduction

Hollow hydrogel microcapsules provide semi-permeable compartments that retain cells while allowing molecular exchange, supporting diverse applications from clonal expansion and secretion-based screening to standardized spheroid culture in microbiology, drug discovery, and cell biology (1–8). Microfluidic methods can produce highly uniform capsules at scale, but most implementations rely on bulky syringe pumps or pressure controllers (9–13), limiting adoption in confined workspaces such as anaerobic chambers for gut microbiota studies and biosafety cabinets for viral research (10-13).

To address this constraint, here we developed a minimal-instrument workflow to generate uniform, cell-laden hollow microcapsules at millions scale, which replaces active pumping and pressure control with a sealed-bottle negative-pressure source and vortex-based particle-templated emulsification (PTE). The entire setup fits within 30 cm × 30 cm and is thus compatible with space-restricted environments such as an anaerobic chamber. We quantify throughput, uniformity, and batch yield, and then demonstrate fully in□chamber operation by encapsulating mouse gut bacteria and tracking population-level proliferation over four days. Together, these results establish a space-efficient route to high-throughput production of uniform, cell-laden hollow agarose microcapsules, positioning the platform for single-cell and clonal analyses in constrained environments.

## Materials and methods

### Preparation of hydrogel solution

#### Fluorescent alginate solution

AF647-labeled alginate was prepared by conjugating Alexa Fluor 647 Hydrazide (Invitrogen, A20502) to sodium alginate (Kimica, IL-6) using N-ethyl-N’-(3-dimethylaminopropyl)carbodiimide hydrochloride (EDC, Merck, E7750) in MES (Sigma, M8250) buffer (pH 5.5) following the protocol for fluorescently-labeled alginate described elsewhere (8). We purified the product by isopropanol (Wako, 166-04836) precipitation and lyophilization before use.

For alginate core preparation, we first prepared two separate alginate solutions. A 2% (w/v) sodium alginate solution was made by dissolving sodium alginate in D-PBS(-) (Wako, 045-29795), and a 2% (w/v) CB-alginate solution was prepared similarly. These two solutions were then mixed at a 9:1 volume ratio. Separately, we prepared an EDTA/Ca buffer (50 mM EDTA, 50 mM CaCl_2_) by diluting 1 M CaCl_2_ (Sigma, 21115-25L) and 0.5 M EDTA (pH 8.0, Thermo Fisher Scientific, AM9260G) in Milli-Q water, adjusting the final pH to 7.2 with 1 M NaOH (Fujifilm, 190-02171). Finally, we mixed the combined alginate solution with the EDTA/Ca buffer at a 1:1 ratio to obtain final concentrations of 1% alginate, 25 mM EDTA, and 25 mM CaCl_2_. The preparation of the alginate mixture for bacterial encapsulation is described in the “Bacteria isolation, encapsulation, and culture” section.

#### Fluorescent agarose solution

Detailed protocols for the preparation of FITC-labeled agarose are described elsewhere (8). Briefly, fluorescein isothiocyanate isomer-I (FITC, Wako, F007) was conjugated to ultra-low gelling temperature agarose (Sigma, A5030-5G) in Dimethyl Sulfoxide (Super Dehydrated DMSO, Wako, 048-32811), followed by repeated centrifugation to remove unreacted FITC, then lyophilized and reconstituted as a 1% solution with Milli-Q water. We prepared an agarose solution for the experiment by dissolving 1.5% (w/v) agarose in an alginate buffer consisting of 10 mM Tris (pH 8.0), 137 mM NaCl, 2.7 mM KCl, and 1.8 mM CaCl_2_, and then mixed it with a 1% FITC-agarose solution in a 97:3 ratio.

### Negative pressure-driven droplet generation

We constructed a negative pressure source using a plastic bottle (Corning, 431531). We equipped the bottle with a pressure gauge (SMC, GZ46-K1K-01) on its sidewall for real-time monitoring and created an outlet by punching a hole in the bottle cap to insert an ETFE tube (OSAKA CHEMICAL, T-082-10). To the end of the ETFE tube, we connected a PC 2-way luer stop cock with male luer hub to male luer lock ring (AKS2FMLW) using a ferrule ring (IDEX, P-250X) and PEEK nut (IDEX, LT-115X) to ensure a secure, leak-proof connection for pressure control. To make the system airtight, we sealed all connection points with adhesive (CEMEDINE Super X2).

We designed microfluidic channels using AutoCAD and fabricated the device with polydimethylsiloxane (PDMS, SILPOT184, Dow) using standard soft lithography techniques. The device contained two inlets: one for alginate solution (or bacterial suspension) and another for oil. The channel geometry is illustrated in Fig. S3.

To generate droplets, we first loaded the alginate solution into a 200 µL pipette tip and connected it to the device’s alginate inlet. Concurrently, we loaded Automated Droplet Generation Oil for EvaGreen (Bio-Rad, 1864112) into a 1 mL pipette tip and connected it to the oil inlet (Fig. S3). The device outlet was connected via PEEK tubing (Nirei Industry, NPK-008) to a 2 mL Safe-Lock tube (Eppendorf, 0030123620) equipped with a septum to maintain an airtight seal. We then connected a 30 mL syringe (Terumo, SS-30LZ) to the ETFE tube from the plastic bottle via a valve and withdrew air until the internal pressure of the bottle reached -20 kPa, at which point we closed the valve to seal the pressure source. Finally, we connected the 2 mL tube to the valve using another PEEK tube and opened the valve to apply the -20 kPa pressure to the microfluidic device, initiating droplet generation.

We calculated the throughput of droplet generation by recording videos at 7,000 frames per second using a high-speed camera (DITECT, HAS-EX). We manually counted generated droplets from 1,000 consecutive frames, and the production rate was calculated by dividing the total count by the corresponding time.

After completing droplet generation, we solidified alginate by lowering the pH. The addition of 0.1% (v/v) acetic acid to the oil phase lowered the pH, releasing calcium from EDTA and inducing alginate crosslinking. Alginate beads were recovered by breaking the emulsion with HFE-7200 (3M) containing 20% (v/v) 1H,1H,2H,2H-perfluoro-1-octanol (PFO, Wako, 324-90642), followed by centrifugation for 15 s using a Mini Personal Centrifuge (Tomy, HF-120) and supernatant removal to collect the bead pellet.

### Agarose microcapsule generation by templated emulsification

Equal volumes of alginate beads and an agarose mixture were combined, followed by the addition of Automated Droplet Generation Oil for EvaGreen, which was five times the volume of the bead and agarose mixture. We vortexed the mixture at maximum speed for 1 minute (Biosan, Vortex Mixer V-1 Plus) to generate emulsions. The emulsions were placed on ice for 10 minutes to solidify the agarose layer.

After gelation, we broke the emulsion using the same procedure as for alginate bead recovery. Two-layered beads were washed with D-PBS(-) for generating hollow agarose hydrogel microcapsules without bacterial culture, or with Anaerobic Akkermansia Medium (AAM medium) when culturing mouse gut bacteria within the microcapsules. Both washing procedures dissolved the alginate cores, yielding hollow agarose hydrogel microcapsules.

We calculated the total number of microcapsules using a C-chip hemocytometer (DHC-N01). Microcapsules were suspended in 4 mL of D-PBS(-), and 10 μL of the suspension was loaded into the hemocytometer to calculate microcapsule concentration. The total number of microcapsules was then determined by multiplying the concentration by the total volume of the suspension.

### Bacterial culture

#### Preparation of culture medium

Mouse gut bacteria were cultured in AAM medium, a defined medium optimized for the growth of anaerobic gut bacteria (5). To prepare the medium, we dissolved 18.5 g of Brain Heart Infusion (Difco, 237400), 5 g of Yeast Extract (Sigma Aldrich, 70161-50G), 15 g of Trypticase soy broth (Biokar Diagnostics, BK046HA), 2.5 g of K_2_HPO_4_ (Wako, 164-04295), 0.5 g of Glucose (Wako, 045-31162), 0.4 g of Na_2_CO_3_ (Wako, 195-01582), and 0.5 g of Cysteine hydrochloride (Wako, 031-05273) in 1 L of Milli-Q water. The solution was bubbled with N_2_ gas for 5 minutes, autoclaved, and cooled in a plastic bag infused with N_2_ gas. After cooling, we transferred the medium to an anaerobic chamber (Sheldon, Bactron300) and added 2 mL of 2.5 mg/mL Menadione (Wako, 134-08131), 200 μL of 5 mg/mL Hemin (Wako, 089-10321), and 30 mL of fetal bovine serum (FBS, Sigma, F7524).

#### Bacteria isolation, encapsulation, and culture

We purchased 8-week-old male C57BL/6J mice from either CLEA Japan or Oriental Yeast for all experiments. To isolate the cecal microbiota, we dissected the cecum, incised it, and submerged it in 3 mL of D-PBS(-). The contents were released by gentle agitation to create a bacterial suspension, which was then filtered through a 100 μm cell strainer (Greiner, 542000) to remove large debris. We subsequently adjusted the optical density (OD) of the bacterial suspension to 0.5 at 600 nm after a 100-fold dilution in D-PBS(-). For encapsulation, we created a mixture by combining a 2% alginate solution, the prepared bacterial suspension, a 200 mM Ca-EDTA solution, and D-PBS(-) at a volume ratio of 20:1:5:14, respectively. After completing the three-step encapsulation workflow, we cultured the resulting microcapsules in 10 mL of AAM medium at 37°C within an anaerobic chamber. At designated time points, we collected 1 mL of the microcapsule suspension for imaging.

All animal experiments were performed in accordance with procedures approved by the Research Center for Advanced Science and Technology, The University of Tokyo, and conformed to the university’s guidelines for the care and use of animals.

### Fluorescence imaging

To quantify bacterial proliferation, we stained microcapsule suspensions with DAPI (Thermo Fisher Scientific, D3571) immediately before imaging at each time point. We collected 1 mL of the suspension and washed it twice with D-PBS(-) by centrifuging at 300*g* for 2 min and removing the supernatant. The washed microcapsules were then resuspended in 400 µL of a 25 µg/mL DAPI solution in D-PBS(-) and incubated for 15 min at room temperature in the dark. Following incubation, we washed the microcapsules once with 1 mL of D-PBS(-) (300*g*, 2 min) to remove excess dye before imaging.

We performed imaging using the EVOS M7000 Imaging System (Thermo Fisher Scientific, C10228). Images were recorded as 12-bit grayscale TIFF files with a resolution of 2048 × 1536 pixels. Light intensity, exposure time, and gain were kept consistent across similar experiments.

### Image analysis

Fluorescent images were analyzed using custom Python scripts based on SciPy 1.9.3, scikit-image 0.24.1, OpenCV 4.9.0, matplotlib 3.7.1, and NumPy 1.26.4.

#### Analysis of core alginate gels and agarose microcapsules

We developed an image analysis pipeline to identify and measure alginate cores and agarose microcapsules from TIFF images. First, we preprocessed raw images by applying a Gaussian filter (sigma = 1) to reduce noise and then binarized them using Otsu’s thresholding. To remove artifacts and fill holes in the binary masks, we then applied specific sequences of dilation and erosion. To ensure precise segmentation, we selected kernel sizes after inspecting the merged mask and phase contrast image. For alginate cores, we applied morphological dilation followed by erosion with a 3×3 kernel; for agarose microcapsules, we applied erosion with a 5×5 kernel. After these refinements, we identified individual objects by labeling the connected components. We then calculated the diameter of each object from its area, assuming a circular shape. Finally, we filtered these objects based on their geometric properties to exclude incorrectly segmented items, such as small debris and large aggregates. We retained alginate cores with an area between 100 and 10,000 pixels and a circularity of 0.90 or greater. Similarly, we retained agarose microcapsules with an area between 5,000 and 20,000 pixels and a circularity of 0.90 or greater. Circularity was calculated using the formula: 4π * area / (perimeter^2^).

#### Quantification of bacterial proliferation in microcapsules

To quantify the DAPI signals, we first segmented the agarose microcapsules from the corresponding FITC-agarose images. The preprocessing steps, including Gaussian filtering (sigma = 1) and Otsu’s thresholding, were performed as described above. We then refined the binary masks with erosion followed by dilation, both using a 5×5 square kernel; these morphological operations were applied to remove small artifacts. Touching microcapsules were separated using a watershed algorithm with a 3×3 footprint. Finally, similar to the previous analysis, we filtered the segmented objects by area, retaining only those between 7,000 and 10,000 pixels.

To quantify bacterial proliferation, we first established a detection threshold to distinguish between microcapsules containing bacteria and those that are empty. Control microcapsules without bacteria were stained with DAPI and segmented; the mean DAPI intensity was then measured for each capsule to determine the background DAPI signal. We performed this analysis across three independent experiments and calculated two statistical metrics: the 99th percentile and the mean plus three standard deviations of the control signals. To ensure a conservative detection and minimize false positives, we adopted the maximum of these two values as our final threshold (Fig. S2). The proportion of capsules with detectable bacteria was then calculated for each time point by dividing the number of capsules with a mean DAPI intensity exceeding this threshold by the total number of segmented capsules.

## Results

### Generation of uniform hollow agarose microcapsules using a space-efficient microfluidic setup

We implemented a three-step process (Fig. 1A). In Step 1, alginate droplets containing cells (AF647-labeled for visualization) are formed inside a microfluidic flow focusing device by applying −20 kPa via a sealed bottle; 0.1% (v/v) acetic acid in the oil lowers pH to release Ca^2+^ from EDTA and subsequently crosslink alginate. In Step 2, the alginate beads are used as templates to form the hollow microcapsules of agarose (15) (FITC-labeled for visualization) with particle-templated emulsification (PTE), wherein the alginate beads are resuspended in an agarose solution, emulsified by vortexing (14), and cooled to form core–shell hydrogel beads. In Step 3, exposing the two-layer hydrogel beads to the Ca^2+^-free buffer causes dissolution of the alginate cores, yielding hollow agarose microcapsules.

**Fig. 1.**
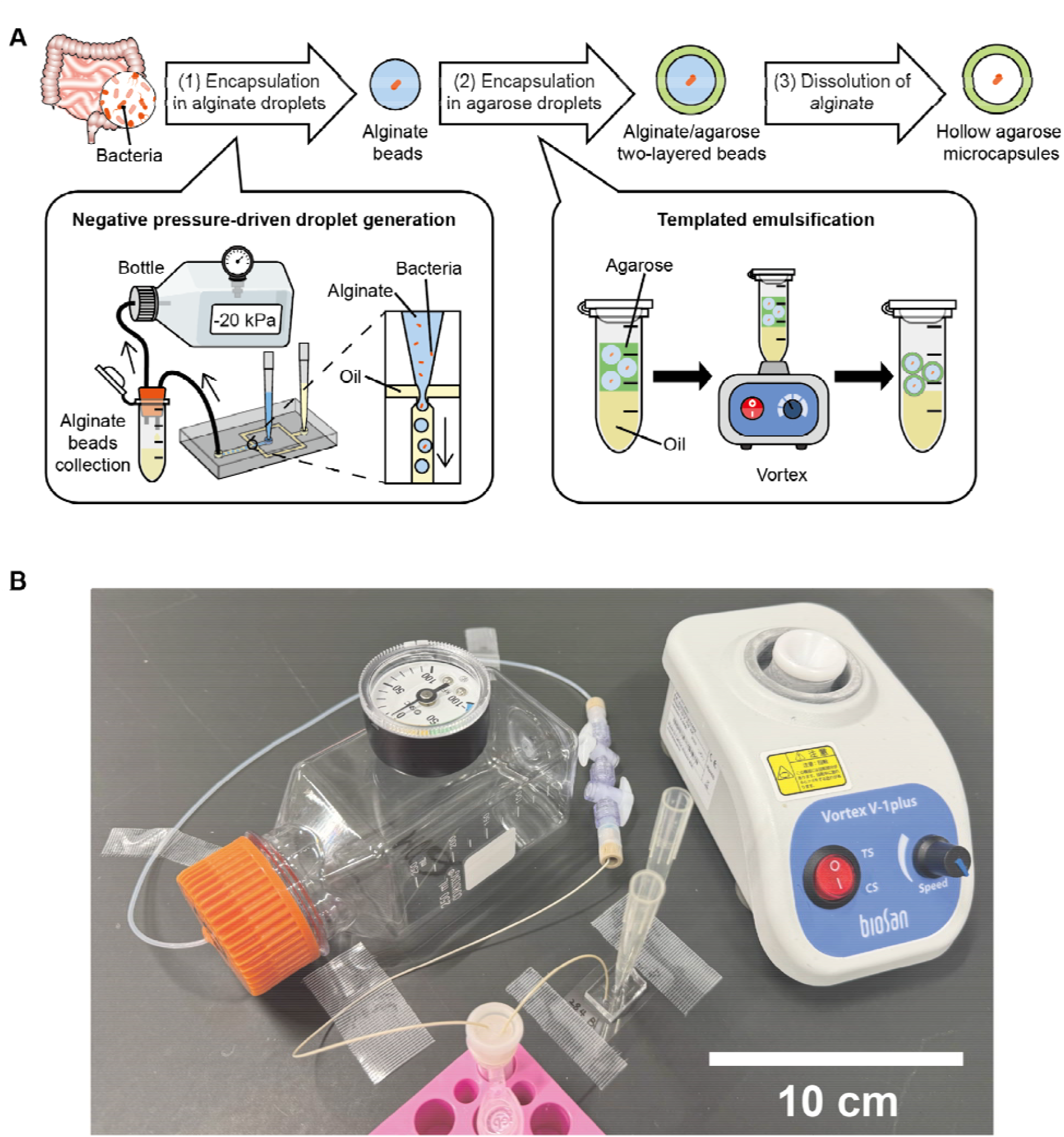
Workflow for the generation of uniform-sized hollow microcapsules with a space-efficient setup. (A) Three-step process for producing hollow agarose microcapsules. (i) Pump-free, −20 kPa negative-pressure droplet formation of alginate; gelation is triggered by pH decrease to release Ca^2+^ from EDTA, followed by de-emulsification to recover alginate beads. (ii) Particle-templated emulsification: alginate beads are encapsulated in agarose droplets using a vortex mixer; cooling solidifies the agarose, and de-emulsification recovers two-layer hydrogel beads. (iii) Core-alginate dissolution in Ca^2+^-free medium generates hollow agarose microcapsules. (B) Photograph of the complete setup comprising a sealed bottle, a disposable microfluidic chip, and a vortex mixer, for the microcapsule generation.

High-speed imaging at 10-min intervals over 100 min showed stable droplet production in Step 1 at a throughput of 491 ± 2 drops/s (mean ± SD across three independent experiments; Figs. 2B and S1). A custom Python-based automated fluorescence image analysis confirms that the solid alginate core beads are uniform with 55.9 ± 0.3 µm diameter and a coefficient of variation (CV) of 3.6 ± 0.7% (mean ± SD, N = 3; Fig. 2C). After the Step 3, a similar Python-based analysis confirms that the hollow-core agarose capsules are uniform with 68.1 ± 0.3 µm diameter and a CV of 3.9 ± 0.4% (mean ± SD, N = 3; Fig. 2D). The sizes of cores and shells imply an average shell thickness ∼6.1 µm. The total production yield was estimated by multiplying the final capsule concentration by the suspension volume is 2.26 × 10^6^ ± 9.17 × 10^4^ capsules per batch (mean ± SD, N = 3; Fig. 2E).

**Fig. 2.**
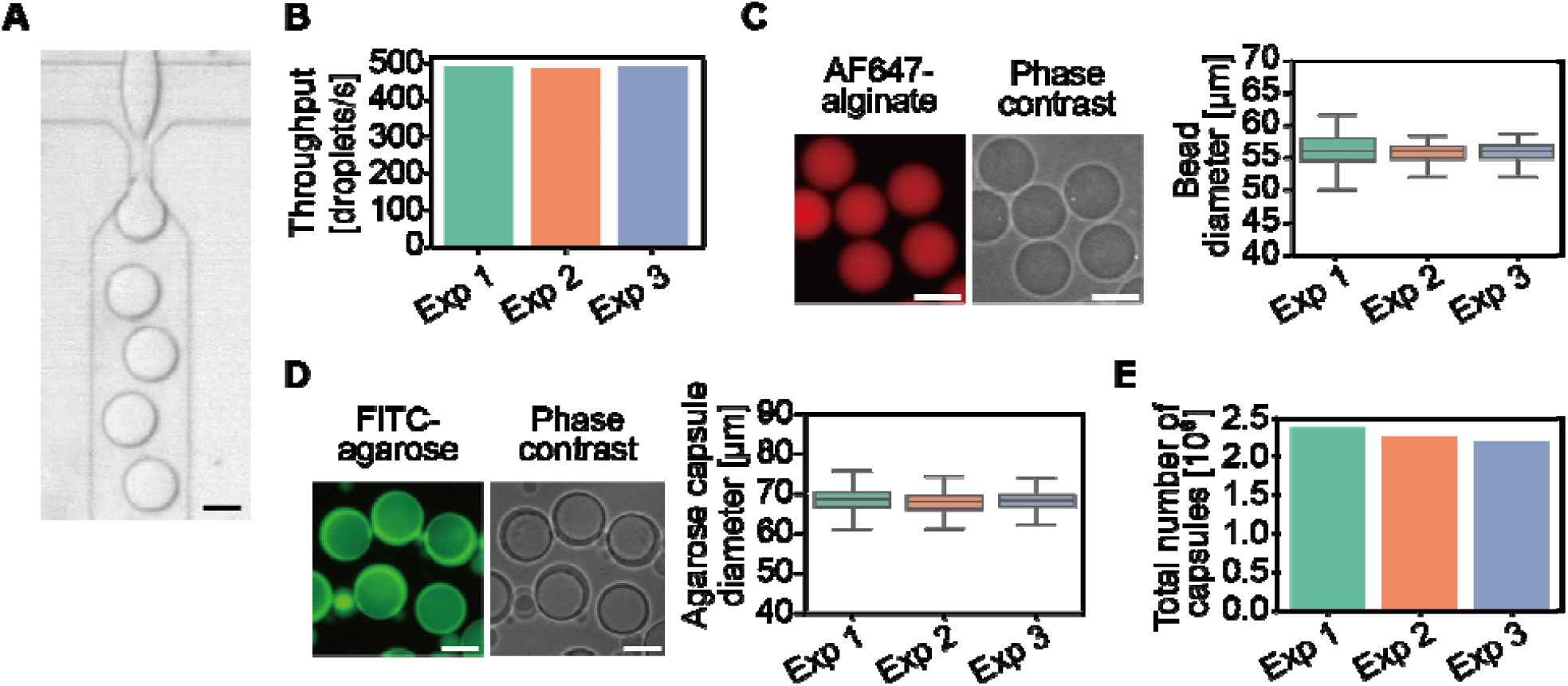
Large-scale production of uniform agarose microcapsules using a space-efficient setup. (A) Representative brightfield image of alginate droplet generation within the microfluidic channel. Scale bar: 50 μm. (B) Throughput over 100□minutes sampled at 10-minute intervals across three experiments. (C) Representative fluorescence and phase-contrast images of AF647-labeled alginate beads (left) and box plots of their diameter distributions (right). Sample sizes were 1073, 1038, and 932 beads, respectively. Scale bars: 50 µm. (D) Representative fluorescence and phase-contrast images of FITC-labeled agarose microcapsules (left) and box plots of their outer diameter distributions (right). Sample sizes were 1990, 1291, and 926 capsules, respectively. Scale bars: 50 µm. (E) Total yield of microcapsules per batch from three independent experiments, calculated by multiplying the microcapsule suspension concentration by its volume. Colors and labels (Exp 1-3) are consistent experimental replicates across panels B-E.

### In-chamber encapsulation supports anaerobic bacterial proliferation

To test operation in a confined anaerobic environment, we performed all steps inside a chamber, including the generation of droplets with mouse gut microbiota; PTE and hollow-core capsule formation; and bacterial culture. Bacteria encapsulated in FITC-agarose microcapsules were stained with DAPI at the indicated time points and imaged to quantify population□level proliferation of bacteria (Fig. 3A).

**Fig. 3.**
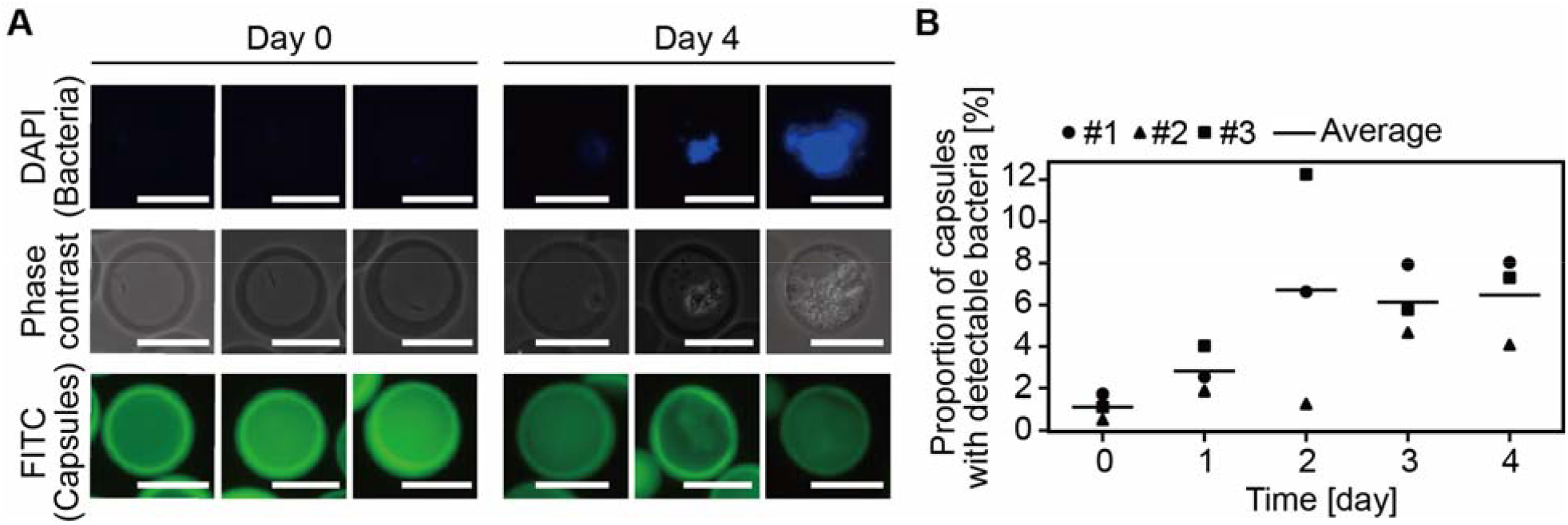
Microcapsules support the long-term culture of mouse gut bacteria in an anaerobic chamber. (A) Representative fluorescence and phase-contrast images of mouse gut bacteria cultured inside agarose microcapsules for four days under anaerobic conditions. FITC-agarose capsules (green) with DAPI-stained bacteria (blue). Scale bars: 50 μm. (B) Time course of bacteria-containing (DAPI-positive) capsule fraction over four days of anaerobic culture. The positivity threshold equals the maximum of the 99th percentile and mean + 3 SD from bacteria-free controls (Fig. S2). Individual data points represent results from three independent experiments, and the lines indicate the mean.

In a custom image analysis developed, we first established a detection threshold to distinguish microcapsules containing bacteria from bacteria-free DAPI-stained controls (maximum of the 99th percentile and mean + 3 SD across three independent experiments; Fig. S2; final threshold = 78.5 intensity units). We then identified bacteria-containing microcapsules when the mean DAPI intensity inside each exceeded the threshold. The fraction of DAPI-positive microcapsules increased from 1.1 ± 0.6% at day 0 to 6.7 ± 5.5% at day 2 and remained 6.5 ± 2.1% at day 4 (mean□± SD, N = 3; Fig. 3B). In these experiments, we encapsulated bacterial cells at a low concentration such that most microcapsules contained zero or one bacterium based on Poisson distribution. These results confirm that the compact workflow supports encapsulation and growth of anaerobic bacteria entirely within a space-restricted chamber.

## Discussion

This study demonstrated that negative-pressure droplet generation coupled with PTE enables space-efficient, high-throughput production of uniform hollow microcapsules using only a sealed bottle and a vortex mixer. The system sustains ∼500 drops per second, achieves a consistent microcapsule size (68.1 ± μm diameter, N = 3) with CV < 4%, and yields ∼2.3 × 10^6^ capsules per batch. Importantly, unlike bulky instruments needed for conventional droplet microfluidics, the workflow operates completely inside an anaerobic chamber, where we observed a growth of mouse gut bacteria inside the capsules over four days.

Our method enables in-capsule, single-clone analysis in anaerobic chambers and biosafety cabinets, thereby accelerating studies of anaerobic and pathogenic bacteria. Compared with double-emulsion approaches (16), agarose microcapsules may offer greater mechanical robustness and simplified handling while avoiding high-pressure gas or multi-pump setups. Encapsulating rare or slow-growing anaerobes at single-clone resolution is thereby made more accessible in constrained environments.

However, several limitations require consideration. Residual unencapsulated bacteria can proliferate outside capsules. Potential post-encapsulation solutions include density-based separation, graded filtration, or flow sorting for purifying microcapsules. Additionally, rapid bacterial growth occasionally caused microcapsule rupture, which could be addressed by thicker capsule walls or selective removal of overgrown capsules.

## Conclusion

By replacing bulky active pumps and pressure controllers with a sealed-bottle negative-pressure source and PTE, we deliver a minimal-instrument workflow that produces millions of uniform hollow agarose microcapsules within a compact footprint and functions entirely in anaerobic chambers. This accessible and robust platform enables single-cell research and clonal expansions in confined spaces and lays a foundation for integrating in-chamber sorting and screening, accelerating discoveries in microbiome and infectious disease research.

## Author contributions

Conceptualization, S.O. and K.H.; Methodology, B.X., S.K., H.K., K.H., and S.O.; Software, K.H. and B.X.; Validation, B.X.; Formal Analysis, B.X. and K.H.; Investigation, B.X. and S.K.; Resources, B.X., H.K., and K.H.; Data Curation, B.X.; Writing – Original Draft, B.X., K.H., and S.O.; Writing –Review & Editing, B.X., S.K., H.K., K.H., and S.O.; Visualization, B.X.; Supervision, S.O. and K.H.; Project Administration, K.H. and S.O.; Funding Acquisition, S.O. and K.H.

## Data availability

Source data for all figures, including the data underlying Figs. S1 to S3 are available from the corresponding author upon reasonable request. The codes are available at https://github.com/solabtokyo-org/.

## Acknowledgements

We thank all members of the Networked Biophotonics Group for their fruitful discussions. A part of Fig. 1 was created with BioRender.com.

This work was supported by JST CREST grant number JPMJCR19H1 and JPMJCR23B6 (to S.O.), JST GteX grant number JPMJGX23B1 (to S.O.), JST as part of Adopting Sustainable Partnerships for Innovative Research Ecosystem (ASPIRE) grant numbers JPMJAP2416 and JPMJAP24B5 (to S.O.), JSPS KAKENHI grant number JP25K22885 (to K.H.), and Japan Agency for Medical Research and Development (AMED) grant numbers JP23tk0124003 (to S.O.) and JP22gm6710008 (to K.H.). This work was also supported by Takeda Science Foundation (to S.O. and K.H.), The Cell Science Research Foundation (to K.H.), Institute for Fermentation (to K.H.), and Ono Medical Research Foundation (to K.H.).

