## Supplemental figures for "Space-efficient method for high-throughput generation of uniform cell-laden hollow agarose microcapsules"

†Co-first authors

*Co-corresponding authors

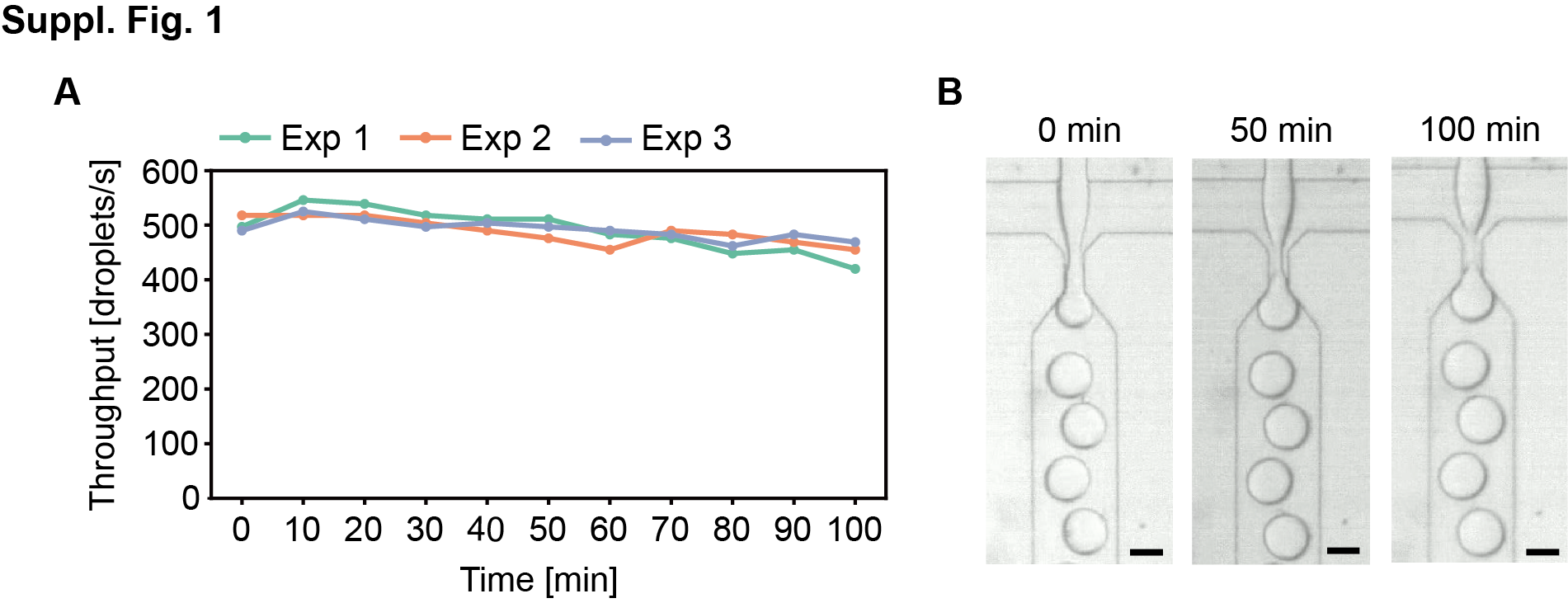
**Figure S1 Temporal changes in the throughput of alginate droplet generation**

(A) Throughput of alginate droplet generation over time, measured at 10-minute intervals over a 100-minute period across three independent experiments. The colors match those in Figure 2.

(B) Representative bright-field images of alginate droplets generated inside microfluidic channels under three different time points. Scale bars: 50 μm.

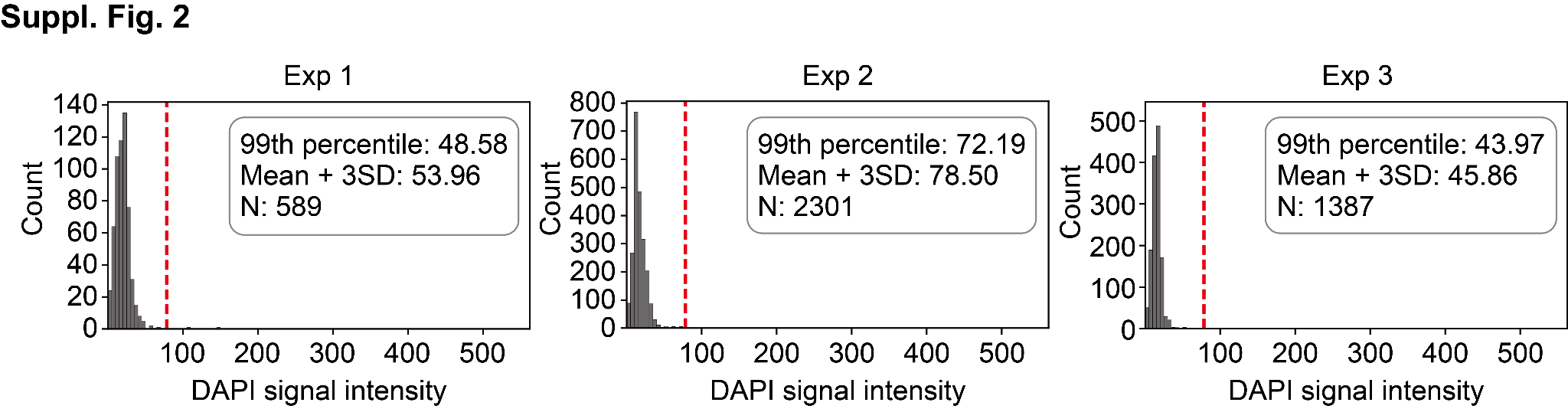

**Figure S2 Statistical determination of DAPI intensity threshold from bacteria-free microcapsule distribution analysis**

Distribution of per capsule DAPI signal intensities from bacteria-free microcapsules across three independent experiments. Microcapsules were stained with DAPI and segmented, and the mean DAPI intensity was calculated for each capsule. These values represent the background DAPI signal in the absence of bacteria. For each experiment, the 99th percentile and the mean plus three standard deviations were calculated as candidate thresholds. To ensure a conservative cutoff that minimizes false positives, the maximum value among all candidate thresholds across experiments, 78.5 intensity units, was adopted as the final DAPI intensity threshold to distinguish microcapsules containing bacteria from bacteria-free controls. The red dotted line in each plot indicates this threshold.

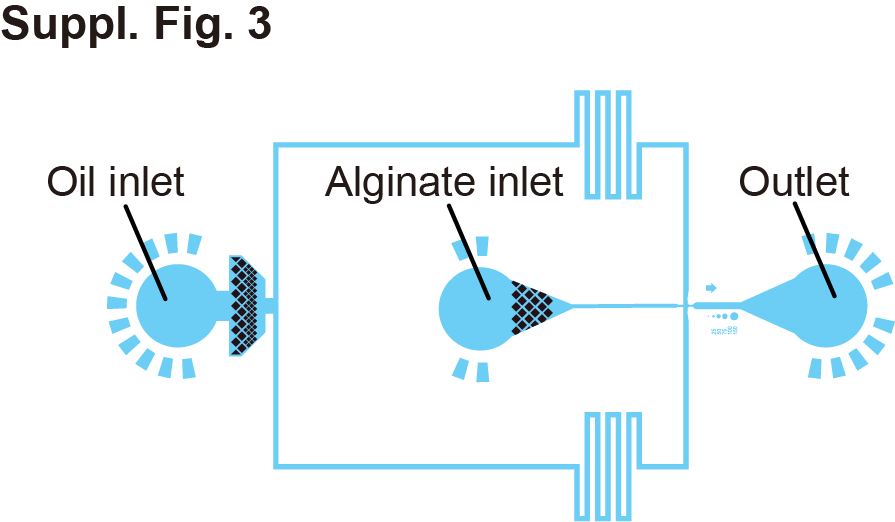

**Figure S3 Design of the microfluidic device for droplet generation**
